# PyMSQ: a python package for fast Mendelian sampling (co)variance and haplotype-based similarity in genomic selection

**DOI:** 10.1101/2025.10.14.682303

**Authors:** Abdulraheem Arome Musa, Norbert Reinsch

## Abstract

**Background:** While genomic selection (GS) boosts rapid genetic gains by leveraging dense marker data for genomic estimated breeding values (GEBVs), prolonged application can reduce haplotype diversity and increase inbreeding. To address these risks, recent research has emphasized Mendelian sampling variance (MSV) and covariance (MSC), which capture within-family (co)variation not fully reflected in GEBVs. In parallel, theoretical advances have introduced a haplotype-based similarity measure that targets shared heterozygous segments, enabling more direct control over haplotype diversity—either standalone or in combination with conventional coancestry-based and genomic relationship matrices.

**Results:** We present PyMSQ, an open-source Python package that implements two key developments: (1) a matrix-based approach for computing MSV and MSC in single-trait, multi-trait, and zygotic contexts, and (2) a haplotype-based similarity metric. By combining this matrix-based framework with optimized scientific libraries, PyMSQ achieves up to 332-fold faster computations than gamevar—a publicly available alternative—while preserving numerical accuracy. Using a Holstein-Friesian dataset, we demonstrate PyMSQ’s effectiveness in deriving MSV and MSC, as well as its novel similarity measure, which complements standard genomic relationship matrices by explicitly quantifying shared heterozygous segments rather than overall allele sharing, thereby providing additional insights for balancing immediate gains with long-term diversity.

**Conclusion:** By facilitating the practical use of MSV, MSC, and a haplotype-based similarity metric, PyMSQ enables breeders and quantitative geneticists to adopt haplotype diversity constraints—whether as a standalone criterion or in synergy with optimal contribution selection. This framework opens new possibilities for preserving key haplotypic segments, ultimately supporting more sustainable genomic selection strategies. PyMSQ is freely available under an MIT License at https://github.com/aromemusa/PyMSQ.

## Background

Genomic selection (GS) relies on genomic prediction, which uses dense marker data and predictive models to estimate breeding values more accurately than traditional pedigree-based methods [1, 2]. Although genomic estimated breeding values (GEBVs) have expedited short-term genetic progress, relying solely on GEBVs—without integrating additional criteria such as inbreeding— may exacerbate inbreeding and erode haplotype diversity over time [3–6]. Preserving genetic variation at the haplotype level is especially important for traits influenced by multiple linked loci, where the loss of crucial segments could irreversibly limit future breeding opportunities.

A key step toward more sustainable breeding decisions involves quantifying within-family genetic variation by way of Mendelian sampling variance (MSV) and covariance (MSC). These metrics require phased genotypes, a marker map, and estimated marker effects. Unlike GEBVs— an individual’s average genetic merit—MSV and MSC capture the range of possible gametic outcomes a parent (or mating pair) could produce [7–15]. Identifying parents whose gametes exhibit high variability for one or multiple traits can help breeders “hedge” against narrowing genetic diversity [14, 16–21]. MSV therefore fulfils two complementary roles. First, it supports short-term gain: parents with larger MSV increase the chance that at least a few offspring will outrank their own GEBV, thereby raising the expected within-family response (the rationale behind strategies such as “usefulness-criterion” selection) [12, 13]. Second, it provides the variance component exploited by optimization frameworks—haplotype-similarity constraints[14, 20], coancestry-based optimal contributions [14], genomic-mating [22] and optimal cross-selection schemes [18]—to balance short-term gain with the long-term preservation of diverse haplotypes. However, computing MSV and MSC at scale has historically been computationally challenging, especially for large populations with high-density marker panels. Early simulation-based methods that enumerate or randomly sample parental haplotypes [7–10] become impractical for tens or hundreds of thousands of markers.

Subsequent analytical methods express breeding values in terms of marker effects [11, 23, 24], yet many still require constructing a parent-specific (co)variance matrix with phase indicators for heterozygous loci—an intensive step in large populations with dense marker sets. In contrast, Musa and Reinsch [14] recently proposed an efficient matrix-based representation that sets up parent-specific marker effects (including phase indicators for heterozygous loci) while building only a single population covariance matrix for each chromosome. This approach avoids creating a unique covariance matrix per parent, thereby enabling rapid computation of MSV and MSC under single-trait, multi-trait, and zygotic models. Beyond speed, the same framework yields a haplotype-based similarity measure that complements coancestry or genomic relationship matrices (GRMs) by targeting shared heterozygous segments (see “Haplotype-Based Similarity” below). This measure quantifies the extent to which MSVs of potential parents arise from identical chromosomal segments. By optimizing mate selection to minimize parental similarities, breeders can preserve broader allelic diversity while maintaining expected genetic gain [14, 20].

Despite this theoretical progress, accessible software implementing both the matrix-based MSV/MSC framework and haplotype-based similarity has been limited. Many breeding programs still rely on simulation-heavy or inefficient analytical methods (e.g., gamevar [24]) that do not provide haplotype-level diversity metrics, or that repeat computationally demanding steps for each parent. To bridge this gap, we developed PyMSQ—an open-source Python package that merges the efficient MSV/MSC derivations of Musa and Reinsch [14] with haplotype-based similarity into a single platform. By leveraging optimized scientific libraries, PyMSQ can handle large datasets, multi-trait analyses, and zygotic variance calculations more feasibly than older methods.

In what follows, we detail PyMSQ’s design and implementation, validate its performance on a Holstein-Friesian dataset, and demonstrate how breeders can integrate MSV, MSC, and haplotype-based similarity to maintain diversity while achieving robust genetic gains in modern genomic selection programs.

## Implementation

### Overview

PyMSQ is a Python 3.8+ package designed to compute MSV and MSC, alongside a haplotype-based similarity metric, using the matrix-based framework presented by Musa and Reinsch [14]. By uniting parent-specific phase indicators with a single population covariance matrix ***R***^*c*^ per chromosome, PyMSQ eliminates the need for parent-specific covariance matrices—thus significantly shortening runtime. Built primarily with NumPy and pandas for data manipulation, it also uses Numba to just-in-time (JIT) compile numeric routines, enabling faster processing of large genomic datasets.

Although written in Python, PyMSQ is compatible with R via the reticulate package, ensuring straightforward integration with R-centric breeding pipelines. The following subsections describe PyMSQ’s internal data structures, matrix computations, and core algorithms. Comprehensive package documentation, including examples and user guides, is available at https://github.com/aromemusa/PyMSQ.

### Data inputs and preparation

All genotype data must be fully phased and free of missing calls; PyMSQ neither performs phasing nor imputation. Users should provide genotype arrays (or DataFrames) in which each marker is assigned one of the two parental haplotypes. A genetic map is also required, specifying a chromosome identifier (e.g., 1, 2) for each marker, along with either a genetic distance 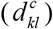 or recombination rate 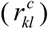 between markers *k* and *l* on a chromosome *c*. These values populate the population-level covariance matrix ***R***^*c*^ for each chromosome *c*, reflecting linkage disequilibrium patterns [14]. Markers on different chromosomes are considered independent.

Finally, marker-effect estimates must be supplied (commonly from a genomic prediction model like genomic best linear unbiased prediction), ensuring their order and dimension exactly match the SNP order of the phased genotype data. In a single-trait setting, PyMSQ expects a vector ***m***^*c*^ ∈ ℝ^*K*^ for each chromosome *c*. For multiple traits, it requires a matrix ***M***^*c*^ ∈ ℝ^*T*×*K*^, where *T* is the number of traits. PyMSQ terminates if it detects inconsistencies in marker order or shape among these datasets, prompting users to correct input files.

### Construction of the population covariance matrix

Musa and Reinsch [14] emphasize constructing a single population covariance matrix ***R***^*c*^ ∈ ℝ ^*K*×*K*^ per chromosome *c*, thereby avoiding parent-specific covariance overhead. The off-diagonal entries 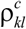 encode the expected recombination-adjusted linkage between markers *k* and *l*. PyMSQ supports two mapping functions:

1. Haldane’s function (assuming no interference) [11, 25]:

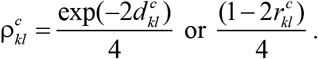
2. Linear approximation (with 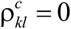 if 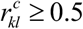 (or 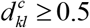 Morgans) [23, 24]:

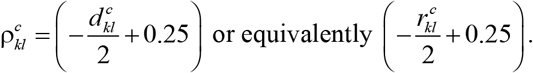

In both models, the diagonal elements of ***R***^*c*^ are 0.25 [11, 14], reflecting the variance contribution of a fully heterozygous locus. Internally, PyMSQ stores each ***R***^*c*^ in a dictionary or list for quick access.

### Parent-specific marker effects

Where earlier methods often built a phase-adjusted matrix 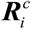 for each parent *i* [11, 23, 24], PyMSQ encodes phase indicators directly into marker-effect vectors (or matrices), following Musa and Reinsch [14]. Let ***m***^*c*^ ∈ ℝ ^*K*^ be the additive marker-effect vector for chromosome *c*. For parent *i*, PyMSQ constructs a vector 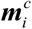 by multiplying each entry in ***m***^*c*^ by a phase indicator *δ*_*ik*_ :

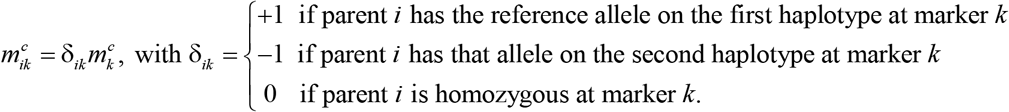

This marker-effect adjustment eliminates the need for parent-specific covariance matrices and smoothly extends to multi-trait contexts by applying *δ*_*ik*_ to each rows of ***M***^*c*^ ∈ ℝ ^*T*×*K*^.

### Computing MSV and MSC for gametes

Once ***R***^*c*^ and 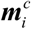 are established, the MSV of parent *i* (single trait) becomes:

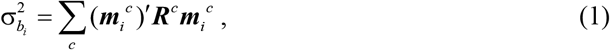

while in multi-trait settings, each chromosome’s marker-effect matrix 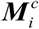 yields:

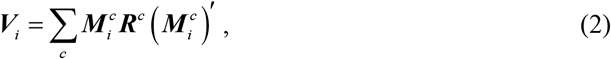

where ***V***_*i*_ contains MSVs on the diagonal and MSCs off-diagonal [14]. An aggregate genotype is computed via ***a****′****V***_*i*_***a***, given a vector of index weight ***a*** ∈ ℝ^*T* ×1^.

### Computing MSV and MSC for zygotes (parent-pair)

When considering a parent pair *ij*, PyMSQ sums each parent’s gametic variance to derive the (co)variance of their potential offspring [11]:

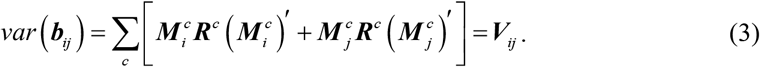

In single-trait mode, this reduces to 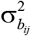. This enables direct comparisons of within-family (co)variation for multiple mating pairs under single- or multi-trait models. The aggregated genotype variance for pair *ij* is ***a****′****V***_*ij*_ ***a***, and PyMSQ outputs each trait’s MSV, the aggregate MSV, and any inter-trait covariances.

### Haplotype-Based Similarity

PyMSQ implements the haplotype-based similarity measure from Musa and Reinsch [14], explicitly targeting shared heterozygous segments among parents. Specifically, the similarity measure quantifies the expected covariance between the additive genetic values of gametes (Mendelian sampling terms) produced by pairs of parents. In single-trait scenarios, the similarity *s*_*ij*_ between parents *i* and *j* is:

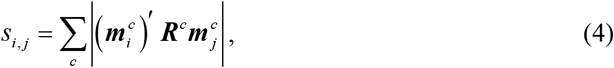

where the absolute value maintains invariance to haplotype order. A similarity matrix ***S*** ∈ ℝ ^*N*×*N*^ for *N* parents can be set up such that a parent’s self-similarity equals its MSV, i.e., 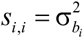. Marker-effect values in 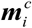 and 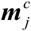 are set to zero for loci homozygous in both parents, as these loci do not generate MSV and hence do not contribute to similarity. Consequently, high similarity values identify parent pairs whose MSVs originate from the same chromosomal segments, whereas low similarity values indicate parents with MSVs arising from distinct chromosomal regions. Breeding schemes can thus optimize mate selection by minimizing average similarity *s*_*ij*_, effectively preserving allelic diversity and moderating long-term inbreeding without compromising immediate genetic gain.

For multi-trait contexts, a user-specified index-weight vector ***a*** yields:

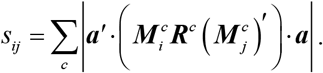

PyMSQ also calculates similarities *s*_*ij*,*uv*_ between zygotes from parent pairs *ij* and *uv* :

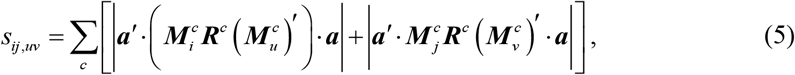

placing the respective MSVs on the diagonal [14]. Here, the row vectors in 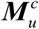and 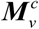 correspond to the trait-specific marker effects of parents *u* and *v* for chromosome *c*.

### Standardized similarity

Because similarity values can range widely, PyMSQ offers a standardized matrix ***K*** derived as:

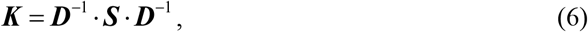

where ***D*** is diagonal, holding each parent’s Mendelian standard deviation 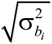. This transforms off-diagonal entries to [0,1] and can help breeders impose constraints on haplotype similarity [14, 20].

### Selection criteria

PyMSQ calculates GEBVs for parent *i* as 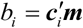, with 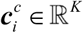 denoting the marker genotype and ***m*** ∈ ℝ ^*K*^ the marker-effect vector derived from genomic prediction models [1]. It then derives an index 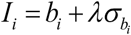, combining GEBV and MSV, where 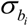 represents the Mendelian standard deviation. The constant *λ* may reflect selection intensity [13] or optimize the probability of producing top-ranking offspring (e.g., 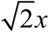, where *x* is the standardized normal truncation point of the selected proportion [12]). Under a zygotic approach, GEBVs and indices for a parent pair *ij* are averaged to explore the joint potential.

### Algorithmic and software optimization

Most core operations in PyMSQ—such as setting up ***R***^*c*^, 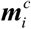 or 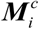 —are vectorized or just-in-time (JIT) compiled using Numba, reducing Python’s overhead in loops and arithmetic on large arrays. For instance, repetitive tasks like 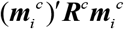 benefit significantly from JIT speedups. The software caches each parent’s phase-adjusted marker effects to avoid recomputing 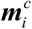 or 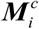 multiple times. It also parallelizes matrix multiplications, leveraging NumPy’s underlying BLAS/LAPACK routines.

### Outputs and integration

PyMSQ returns results primarily as NumPy arrays or pandas DataFrames, facilitating export to .csv or .npy. Because many breeding analyses are R-based, PyMSQ is fully callable from R via reticulate, supporting seamless integration with other R-based genomic pipelines (e.g., rrBLUP, BGLR). By accommodating both Python and R environments, PyMSQ aims to reach a broad community of breeders and geneticists.

We envision that MSV, MSC, and haplotype-based similarity matrices will typically be computed centrally at evaluation centers or large-scale breeding units where phased genotypes and genomic predictions are available. These pre-computed values can then be securely and efficiently distributed (e.g., via routine evaluation reports or secure APIs) to breeding advisors or farms. Thus, direct access to raw genotype data at the farm level is unnecessary. Furthermore, PyMSQ readily accommodates complex multi-trait indices involving numerous traits or sub-indexes, since multi-trait indices are easily represented by single weighted marker-effect vectors, ensuring computational efficiency.

### Data and analysis

Using PyMSQ and a publicly available Holstein-Friesian cattle dataset [14], we derived MSCs and haplotype similarity matrices for the aggregate genotype of gametes, assigning equal index weights of 1. The dataset includes 265 cows from five paternal half-sib families (sizes 32–106), along with a physical map, genotypic data, pedigree information, and phenotypes for three milk traits—pH, fat yield (FY), and protein yield (PY) [26, 27]. Marker effects for this population had been estimated previously [26], and the genotypes were phased into haplotypes (yielding 10,304 markers). Approximate genetic map positions were also established [14]. These data files (map, phased genotypes, marker effects, and pedigree details) are bundled with PyMSQ to ensure full reproducibility.

Following the computation of MSVs and MSCs in PyMSQ, we visualized the results with ggplot2 (v 3.3.3) [28], generating density plots of MSVs and MSCs. We then constructed and examined haplotype similarity matrices using corrplot. We then generated and examined haplotype similarity matrices with corrplot (v 0.84) [29] to assess interrelationships among parental genotypes. Supporting scripts and their outputs are included in Additional File 1.

To compare PyMSQ’s performance with gamevar [24], we ran simulation studies with two data groups, each focused on a single chromosome and ten traits. In the Individuals scenario, we fixed the number of markers and varied the number of individuals to test computational scalability with increasing population size. We generated 20 datasets each containing 1,000 markers but varied the number of individuals. In the Markers scenario, we fixed the number of individuals at 500 while varying marker numbers. Each dataset underwent ten replicates, measuring the computation time (minutes) and peak memory usage (GB) needed to compute MSCs, as well as the time and memory required for haplotype similarity matrices. All computations were conducted on a multi-user Linux server (Intel Xeon Gold 6130, 128 CPU cores at 2.10 GHz, 1536 GB RAM)—a high-memory environment typical of large-scale genomic breeding analyses. Detailed scripts and outputs are provided in Additional File 1.

## Results

### Mendelian variances and trait correlations

Applying PyMSQ to the Holstein-Friesian dataset revealed a wide range of MSVs for the three milk traits: pH, FY, and PY. As illustrated in Figure 1A, MSV distributions for each trait showed coefficients of variation exceeding 90%, underscoring the substantial within-family genetic diversity stemming from heterozygous loci and linkage phase [11, 14]. The FY trait in particular exhibited a bimodal distribution—possibly indicating DGAT1 as a major locus—previously noted to affect milk-fat segregation [8, 11, 23, 30]. Moving beyond single-trait analyses, we derived multi-trait MSCs for each cow, thus quantifying how alleles for pH, FY, and PY co-segregate. The resulting correlations (Figure 1B) revealed that while most cows had moderately positive inter-trait correlations, a subset displayed mildly negative correlations—a reflection of individual differences in linkage phases or pleiotropic effects [11, 23, 31]. These findings highlight opportunities for breeders to strategically select or mate parents to enhance beneficial multi-trait associations or reduce antagonistic linkages.

**Fig. 1:**
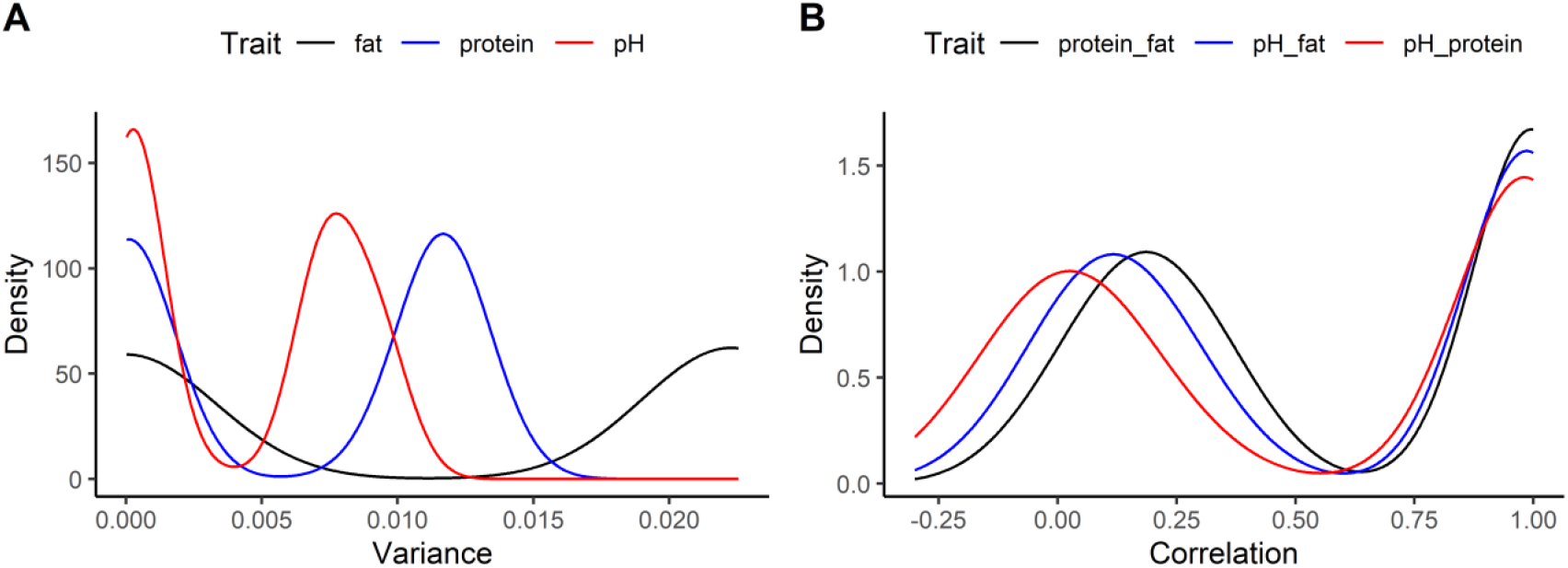
Density plots of Mendelian sampling variances and trait correlations. Panel A displays the variance in fat yield (kg), protein yield (kg), and pH (mol/L). Panel B shows correlations between these traits.

### Haplotype-based similarity and its standardization

We next constructed a haplotype-based similarity matrix for an aggregate genotype combining pH, FY, and PY with equal weights. Diagonal elements in this matrix correspond to each cow’s MSV, and off-diagonals measure the overlap in high-impact heterozygous segments [14]. Figure 2A illustrates that some pairs of cows share extensive haplotype regions, while others remain relatively dissimilar. We also generated a standardized matrix ***K*** (Figure 2B) by dividing off-diagonal entries by the product of each parent’s Mendelian standard deviation, which rescales similarities to a [0,1] interval and accentuates smaller overlaps. This haplotype-level focus complements the GRM by highlighting heterozygous loci relevant to within-family variance, rather than just overall allele sharing [14, 20]. In practical breeding settings, imposing limits on pairwise similarity can retain critical haplotypes, thus promoting the transmission of diverse haplotypes, thereby moderating rather than accelerating long-term inbreeding.

**Fig. 2:**
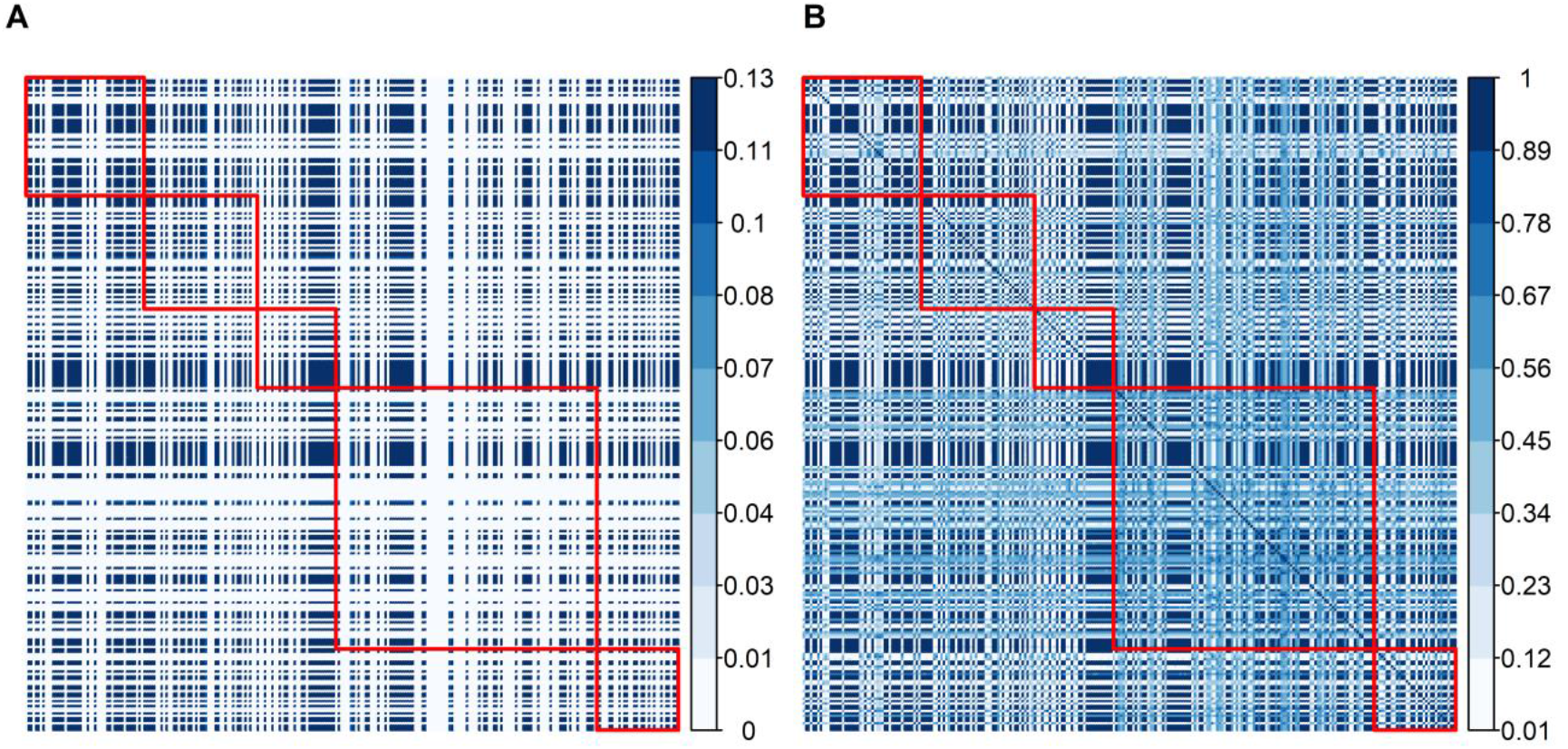
Unstandardized (A) and standardized (B) similarity matrices for the aggregate genotype of 265 Holstein-Friesian cows from five half-sib families (delineated by red lines). In (A), diagonal values represent individual Mendelian sampling variances, and off-diagonal values represent similarity between parent pairs in terms of shared gametic variance. Panel (B) shows the standardized similarity matrix, where values range from 0 (no similarity) to 1 (maximum similarity).

### Comparative benchmarking with gamevar

In the Individuals scenario, the number of markers was fixed at 1,000 while individual counts ranged from 5,000 to 100,000. PyMSQ’s runtime (Figure 3A) increased from about 0.07 min to 1.16 min, whereas gamevar required 1.69 min to 30.80 min—yielding a time ratio (gamevar/PyMSQ) of 23 to 27. Memory usage (Figure 3C) showed the opposite trend: PyMSQ scaled from ~0.55 GB to ~4.97 GB, while gamevar remained near 0.015 GB throughout, reflecting gamevar’s genotype line-streaming approach versus PyMSQ’s in-memory design.

**Fig. 3:**
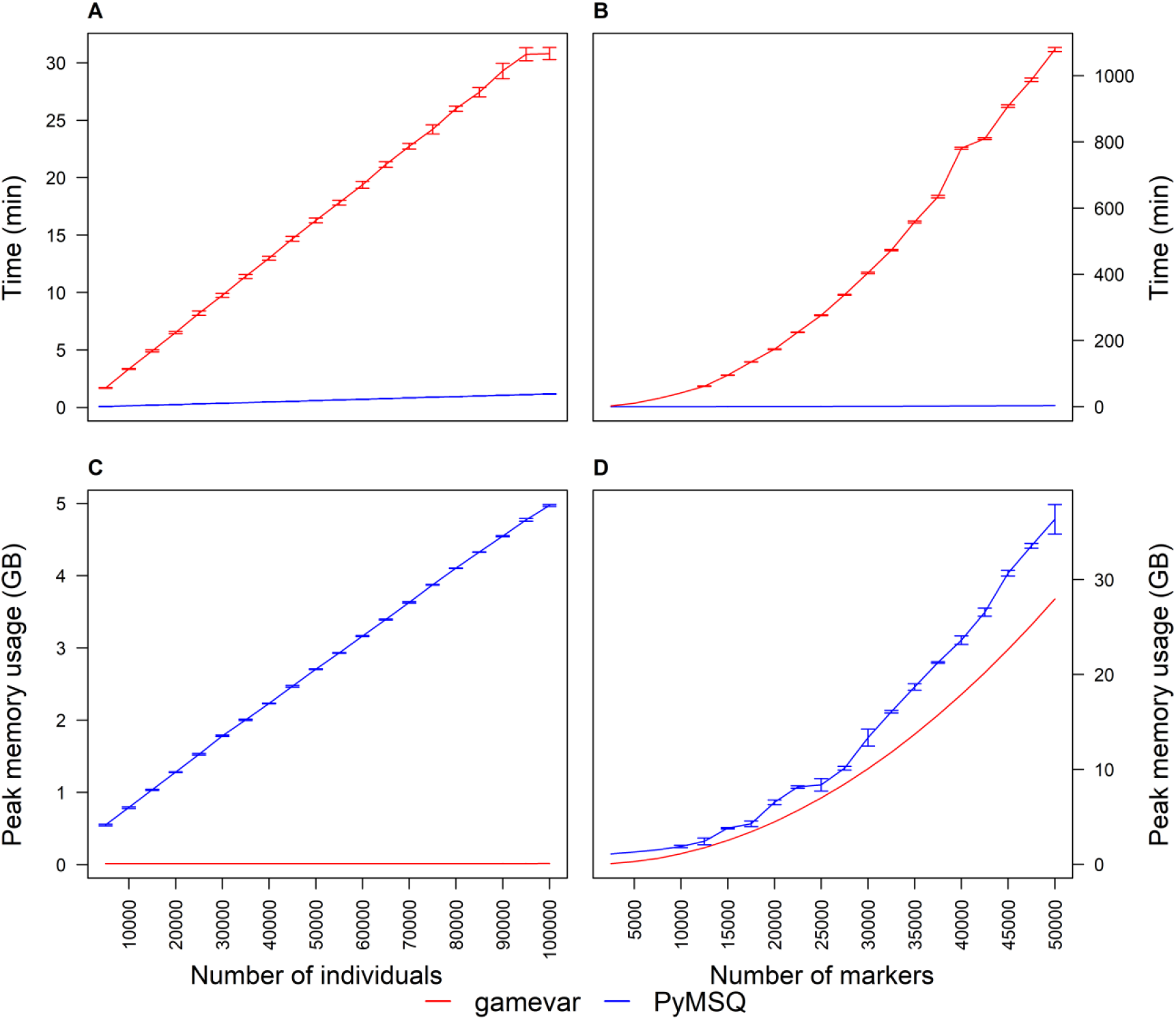
Benchmark plots comparing computation time (Panels A, B) and memory usage (Panels C, D) for PyMSQ and gamevar.

In the Markers scenario (Figure 3B, D), 500 individuals were held constant while marker counts rose from 2,500 to 50,000. PyMSQ’s runtime climbed from 0.03 min to 3.25 min, whereas gamevar soared from 2.44 min to ~1,078.89 min, producing time ratios of up to 332. Gamevar’s memory usage grew from ~0.07 GB to ~27.95 GB, reflecting the quadratic expansion of marker-level arrays. Although PyMSQ’s memory usage also increased (from ~1.10 GB to ~36.35 GB), its matrix-based approach remained markedly faster, particularly for higher marker densities and multi-trait analyses. Overall, PyMSQ provides substantial speed advantages over gamevar, provided sufficient RAM is available.

### Benchmarking of haplotype similarity matrix

We further evaluated PyMSQ’s construction of the haplotype-based similarity matrix, an O(*N*^2^) procedure for *N* parents (Additional File 1, Table 3). In the Individuals scenario (population sizes of 5,000 to 100,000, with 1,000 markers), PyMSQ’s runtime for building this matrix rose from about 0.08 min to 7.93 min, and peak memory from 0.76 GB to nearly 70 GB. Conversely, in the Markers scenario (500 individuals, 2,500–50,000 markers), similarity matrix runtime grew from 0.04 min to 2.8 min, and memory usage climbed from 0.37 GB to ~34.28 GB. Although large servers can handle these demands, breeding programs with constrained resources may choose to subset parents (e.g., based on GEBVs) rather than compute a full *N* × *N* matrix. Nonetheless, PyMSQ’s ability to handle large-scale haplotype similarity computations—whether driven by large *N* or substantial marker densities—enables refined selection and mating strategies that directly target critical heterozygous segments.

### Limitations and future directions

Though PyMSQ is both fast and versatile, its in-memory design can prove demanding for those with restricted hardware. For MSV/MSC calculations, one can process chromosomes sequentially, merging results to reduce memory usage. However, generating a full haplotype similarity matrix for very large *N* may be infeasible unless breeders subset parents based on GEBVs or other criteria. PyMSQ currently focuses on additive genetic models; adding dominance or epistasis would broaden its scope, while further chunk-based or parallel-streaming optimizations could shrink memory overhead. We emphasize that PyMSQ does not replace genomic prediction methods but explicitly serves as a post-processing step applied directly to marker effects derived from genomic prediction methods (Single-Step or Two-Step). PyMSQ explicitly calculates MSV, MSC, and haplotype similarity based exclusively on genomic marker-effect estimates. It does not directly incorporate the additional polygenic (residual additive genetic) component that is explicitly modeled using pedigree or genomic relationships in single-step genomic evaluations. Finally, our tests focused on one cattle population; applying PyMSQ to other species (swine, poultry, crops) should help confirm its broader applicability for sustaining within-family genetic diversity.

## Conclusions

PyMSQ unifies advanced Mendelian (co)variance computation with haplotype-based similarity in a single, open-source platform, enabling faster multi-trait analyses than existing methods. It affords breeders a potent tool to harness within-family variance and safeguard genetic diversity— potentially working in concert with coancestry-based or optimal contribution approaches. Future PyMSQ releases will refine data handling, incorporate non-additive genetic models, and explore multi-species applications, ensuring it remains a practical, high-performance resource for modern genomic selection.

## Supporting information

Additional file 1: Analysis of Holstein-Friesian cattle data and benchmark test

## Availability and requirements

Project name: PyMSQ

Project home page: https://github.com/aromemusa/PyMSQ

Operating systems: Platform independent

Programming language: Python 3.8+

Other requirements: pandas, NumPy, SciPy, and Numba Python libraries

License: MIT license

Any restrictions to use by non-academics: None

All code, documentation, and the Holstein-Friesian dataset used in the examples are available through the GitHub repository. Users may run PyMSQ directly in Python or call it from R via the reticulate package. Contributions—including issue reports, feature requests, or new functionalities—are welcomed via GitHub’s issue tracker.

## Supplementary information

**Supplementary information** accompanies this paper at https://github.com/aromemusa/PyMSQ/blob/main/paper/Additional_file_1.pdf.

**Additional file 1:** Analysis of Holstein-Friesian cattle data and benchmark test

## Abbreviations

FY: fat yield (kg)
GEBV: genomic estimated breeding value
GRM: genomic relationship matrix
GS: genomic selection
JIT: just-in-time (compilation)
MSC: Mendelian sampling covariance
MSV: Mendelian sampling variance
pH: acidity measure (hydrogen-ion concentration)
PY: protein yield (kg)
RAM: random access memory

## Acknowledgments

We thank Nina Melzer and Dörte Wittenburg of FBN Dummerstorf for providing the empirical Holstein-Friesian dataset used in this study.

## Author’s contributions

AAM developed the Python package, analyzed the data, and wrote the manuscript. NR conceptualized and supervised the entire work and revised the manuscript. All of the authors read and approved the final manuscript.

## Availability of data and materials

The code, documentation, illustration, and Holstein-Friesian cattle data are available at https://github.com/aromemusa/PyMSQ.

## Ethics approval and consent to participate

Not applicable.

## Consent for publication

Not applicable.

