## Additional file 1: Analysis of Holstein-Friesian cattle data and benchmark test for "PyMSQ: a python package for fast Mendelian sampling (co)variance and haplotype-based similarity in genomic selection"

### An R script to generate Figs. 1 - 3 and Performance Comparison Tables

08-04-2025

```r
knitr::opts_chunk$set(echo = TRUE)
options(scipen = 999)  # Prevent scientific notation on axes
```

#### Overview

This document loads data, processes it, and generates three figures along with several performance comparison tables:

1. Figure 1: Density plots of Mendelian sampling variances and trait correlations.
2. Figure 2: Similarity matrices and Euclidean clustering of the matrices for the aggregate genotype of milk traits.
3. Figure 3: Benchmark plots comparing computation time and memory usage.
4. Performance Comparison Tables 1 and 2: Summaries from the same benchmark dataset used in Figure 3. These tables report the mean computation time (in minutes) and peak memory usage (in GB) for PyMSQ and gamevar.
5. Performance Table 3: Benchmarking of similarity matrices using PyMSQ.

#### 1. Density Plots

##### Import Packages

We load required libraries

```r
library(ggplot2)      # For creating plots
library(reshape2)     # For reshaping data
library(corrplot)     # For correlation plots
require(RColorBrewer) # For color palettes
library(pheatmap)     # For heatmaps
library(ggpubr)       # For arranging ggplot figures
library(plyr)         # For data manipulation
library(knitr)        # For document generation
library(reticulate)   # For interfacing with Python
```

##### Import Data from PyMSQ

```r
# Replace "pymsq_dev" with the conda environment or virtualenv name.
# use_condaenv("pymsq_dev", required = TRUE)
# py_config()  # confirm your environment is active
```

```r
msq <- import("PyMSQ")
data <- msq$load_package_data()  # now you're set to proceed below


data <- msq$load_package_data()
gmap <- data[["chromosome_data"]]        # genetic map
meff <- data[["marker_effect_data"]]     # marker effects
gmat <- data[["genotype_data"]]          # phased genotype
group <- data[["group_data"]]            # group data
ped <- data[["pedigree_data"]]

# define number of traits and index weight
no_traits <- ncol(meff)                 # 3 traits
index_wt <- c(1, 1, 1)
```

##### Compute Mendelian (co-)variance and correlation using PyMSQ

```r
# Derive population covariance matrix for each chromosome
exp_ldmat <- msq$expldmat(gmap = gmap, group = group, mposunit = "cM")

# Compute Mendelian (co-)variance
msvmsc <- msq$msvarcov(gmat, gmap, meff, exp_ldmat = exp_ldmat, group = group,
                   indwt = index_wt, progress = TRUE)

# Compute Mendelian correlation
mscorr <- msq$msvarcov_corr(msvmsc)
```

##### Prepare and Plot the Density Data

We extract, reshape, and plot the data:

```r
# Extract relevant columns
msv_traits <- data.frame(msvmsc[, c("fat", "protein", "pH")])

# Compute coefficient of variation for each column
round(sapply(msv_traits, function(x) sd(x) / mean(x)) * 100, 2)

mscorr_traits <- data.frame(mscorr[, c("protein_fat", "pH_fat", "pH_protein")])

# Compute mean correlation
round(sapply(mscorr_traits, function(x) mean(x)), 2)

# Compute range of correlations
round(sapply(mscorr_traits, function(x) range(x)), 3)

# Compute percentage of negative values
round(sapply(mscorr_traits, function(x) (sum(x < 0)/265)*100), 1)

# Reshape data
df_msv <- melt(msv_traits)
colnames(df_msv) <- c("Trait", "Variance")
df_mscorr <- melt(mscorr_traits)
colnames(df_mscorr) <- c("Trait", "Correlation")
```

```r
# Create density plots
plot1 <- ggplot(df_msv, aes(Variance, color = Trait)) +
  stat_density(geom = "line", position = "identity") +
  theme_classic() +
  theme(legend.position = "top") +
  scale_color_manual(values = c("black", "blue", "red")) +
  ylab("Density")

plot2 <- ggplot(df_mscorr, aes(Correlation, color = Trait)) +
  stat_density(geom = "line", position = "identity") +
  theme_classic() +
  theme(legend.position = "top") +
  scale_color_manual(values = c("black", "blue", "red")) +
  ylab("Density") +
  scale_x_continuous(limits = c(-0.3, 1), breaks = seq(-0.25, 1, by = 0.25))
```

##### Save and Display Figure 1

We combine the density plots and save them as a TIFF file. We then embed the resulting image:

```
## pdf
##   2
```

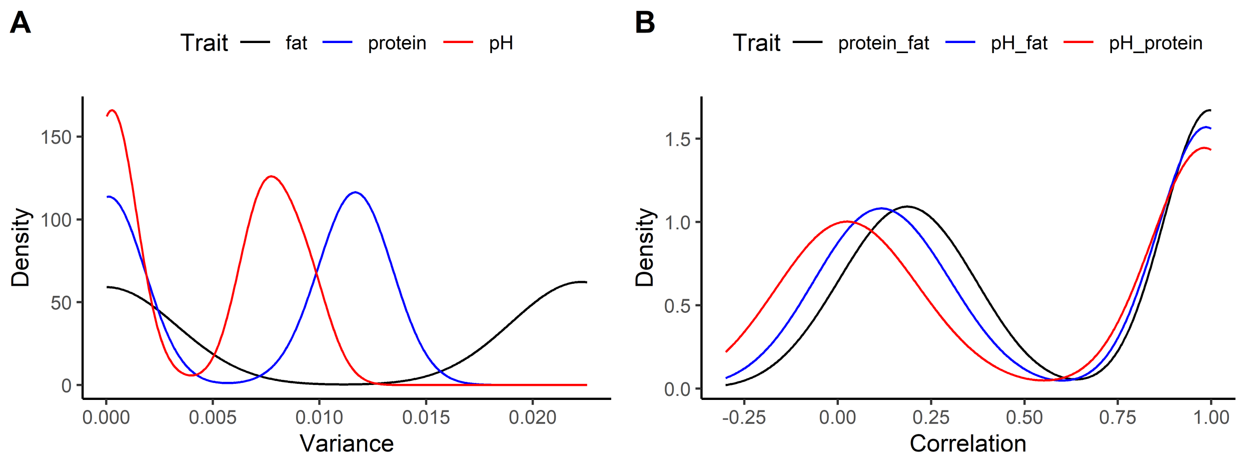


Figure 1: Density plots of Mendelian sampling variances and trait correlations. Panel A displays the variance in fat yield (FY, kg), protein yield (PY, kg), and pH (mol/L). Panel B shows correlations between these traits.

#### 2. Similarity Matrices

We then derive similarity matrices.

```r
# Derive similarity matrix
sim <- msq$simmat(gmat, gmap, meff, group = group, exp_ldmat = exp_ldmat,
              indwt = index_wt, save=TRUE, progress = TRUE)[[1]]
# Standardize the similarity matrix
std_sim <- cov2cor(sim)

# get the range of values from both matrices
```

```r
sim_without_diag <- sim
diag(sim_without_diag) <- NA
range(sim_without_diag, na.rm = TRUE)

std_sim_without_diag <- std_sim
diag(std_sim_without_diag) <- NA
range(std_sim_without_diag, na.rm = TRUE)

# find the minimum value in sim and check the corresponding value in std_sim
minVal <- min(sim_without_diag, na.rm = TRUE)
minPos <- which(sim_without_diag == minVal, arr.ind = TRUE)
correspondingPos <- std_sim[minPos[1], minPos[2]]
```

##### Save and Display Figure 2

```r
# Determine number of individuals within each paternal half-sib family
pedigree <- data[["pedigree_data"]][-1, ]
ped <- as.data.frame(pedigree[,2])
no <- NULL
for (i in 1:length(unique(ped[, 1]))) {
  first <- length(which(ped[, 1] == i))
  if (i == 1) {
    no <- c(no, first)
  } else {
    no <- c(no, first + no[i-1])
  }
}

png("Figure2.png", width = 8, height = 4, units = 'in', res = 700)
par(mfrow = c(1, 2), oma = c(0, 0, 0, 0.1) + 0.1, mar = c(0, 0, 0, 0) + 0.1)
cols <- brewer.pal(9, "Blues")
corrplot(sim, is.corr = FALSE, method = "color", cl.lim = range(sim),
         cl.digits = 1, cl.cex = 0.80, tl.col = "black", tl.pos = "n",
         col = cols, cl.align.text = "c", mar = c(0, 0, 1, 0)) -> p
corrRect(p, c(1, no), col = "red")

corrplot(std_sim, is.corr = FALSE, method = "color", cl.lim = range(std_sim),
         cl.digits = 1, cl.cex = 0.80, tl.col = "black", tl.pos = "n",
         col = cols, cl.align.text = "c", mar = c(0, 0, 1, 0)) -> p
corrRect(p, c(1, no), col = "red")

mtext(expression(bold("A")), side = 3, outer = TRUE, cex = 1, las = 0, line = -1, adj = 0)
mtext(expression(bold("B")), at = 0.52, side = 3, outer = TRUE, cex = 1, las = 0, line = -1)

dev.off()
```

```
## pdf
##   2
```

Now we display Figure 2:

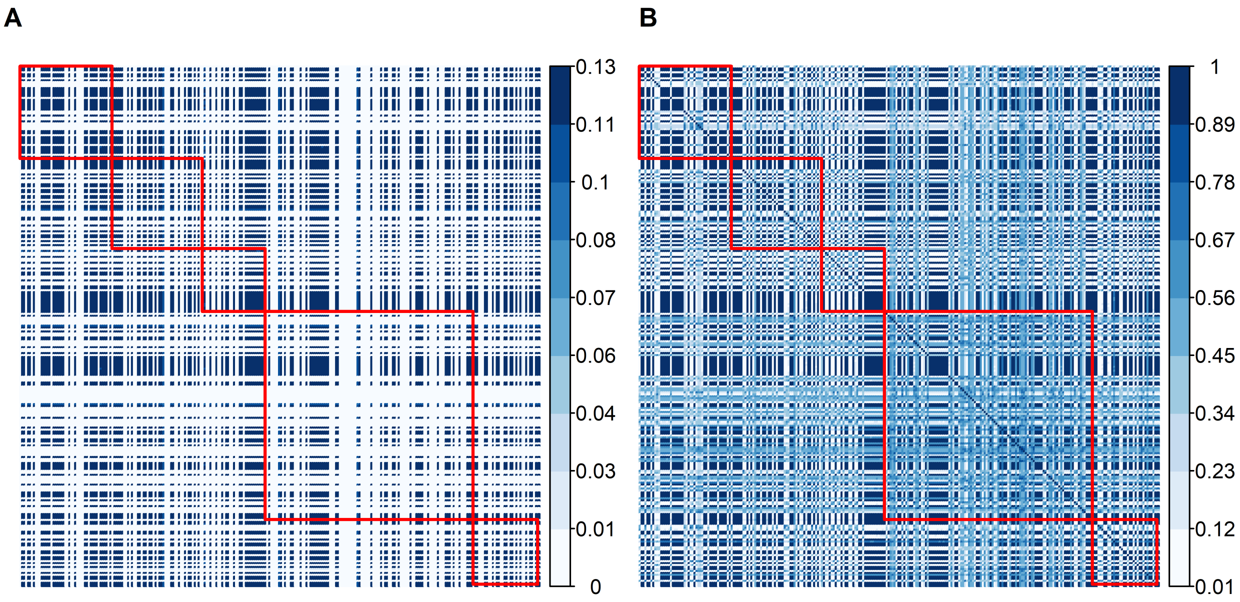


Figure 2: Unstandardized (A) and standardized (B) similarity matrices for the aggregate genotype of some milk traits for 265 cows from five half-sib families (separated by red lines).

#### 3. Benchmark Plots

We then process benchmark data (computation time and memory usage) and generate a 2×2 panel plot.

```
library(dplyr)
library(ggplot2)

# Read csv files
perf_ind <- read.csv("performance_analysis.csv")
perf_mark <- read.csv("performance_analysis_mark.csv")

# Compute computation time (convert seconds to minutes) and its SD
time_ind_summary <- perf_ind %>%
  group_by(no_individuals) %>%
  summarise(
    PyMSQ_mean = mean(time_PyMSQ, na.rm = TRUE) / 60,
    PyMSQ_sd   = sd(time_PyMSQ, na.rm = TRUE) / 60,
    gamevar_mean = mean(time_gamevar, na.rm = TRUE) / 60,
    gamevar_sd   = sd(time_gamevar, na.rm = TRUE) / 60
  )

# Compute peak memory usage (GB) and its SD
mem_ind_summary <- perf_ind %>%
  group_by(no_individuals) %>%
  summarise(
    PyMSQ_mean = mean(peak_memory_usage_PyMSQ, na.rm = TRUE),
    PyMSQ_sd   = sd(peak_memory_usage_PyMSQ, na.rm = TRUE),
    gamevar_mean = mean(peak_memory_usage_gamevar, na.rm = TRUE),
    gamevar_sd   = sd(peak_memory_usage_gamevar, na.rm = TRUE)
  )
```

```
# Compute computation time (in minutes) & its SD (markers)
time_mark_summary <- perf_mark %>%
  group_by(no_markers) %>%
  summarise(
    PyMSQ_mean = mean(time_PyMSQ, na.rm = TRUE) / 60,
    PyMSQ_sd   = sd(time_PyMSQ, na.rm = TRUE) / 60,
    gamevar_mean = mean(time_gamevar, na.rm = TRUE) / 60,
    gamevar_sd   = sd(time_gamevar, na.rm = TRUE) / 60
  )

# Compute peak memory usage (in GB) & its SD (markers)
mem_mark_summary <- perf_mark %>%
  group_by(no_markers) %>%
  summarise(
    PyMSQ_mean = mean(peak_memory_usage_PyMSQ, na.rm = TRUE),
    PyMSQ_sd   = sd(peak_memory_usage_PyMSQ, na.rm = TRUE),
    gamevar_mean = mean(peak_memory_usage_gamevar, na.rm = TRUE),
    gamevar_sd   = sd(peak_memory_usage_gamevar, na.rm = TRUE)
  )
```

##### Save and Display Benchmark Plot (Figure 3)

```
png("Figure3.png", width = 9, height = 8, units = "in", res = 700)
par(mfrow = c(2, 2), oma = c(5,3,1,4.6)+0.1, mar = c(2,1,1,0)+0.1)

# Panel A: Computation Time vs. Individuals
with(time_ind_summary, {
  plot(no_individuals, PyMSQ_mean, type = "l", col = "blue",
       ylim = range(c(PyMSQ_mean - PyMSQ_sd, PyMSQ_mean + PyMSQ_sd,
                      gamevar_mean - gamevar_sd, gamevar_mean + gamevar_sd)),
       xlab = "", ylab = "Time (min)", xaxt = "n", yaxt = "n")
  axis(2, col.axis = "black", las = 2)
  box()
  lines(no_individuals, gamevar_mean, type = "l", col = "red")
  # PyMSQ error bars
  arrows(no_individuals, PyMSQ_mean - PyMSQ_sd,
         no_individuals, PyMSQ_mean + PyMSQ_sd,
         angle = 90, code = 3, col = "blue", length = 0.05)
  # gamevar error bars
  arrows(no_individuals, gamevar_mean - gamevar_sd,
         no_individuals, gamevar_mean + gamevar_sd,
         angle = 90, code = 3, col = "red", length = 0.05)
})
mtext(expression(bold("A")), side = 3, line = 0.5, adj = 0)

# Panel B: Computation Time vs. Markers
with(time_mark_summary, {
  plot(no_markers, PyMSQ_mean, type = "l", col = "blue",
       ylim = range(c(PyMSQ_mean - PyMSQ_sd, PyMSQ_mean + PyMSQ_sd,
                      gamevar_mean - gamevar_sd, gamevar_mean + gamevar_sd)),
       xlab = "Number of Markers", ylab = "", axes = FALSE)
  axis(4, col.axis = "black", las = 2)
  box()
```

```r
  lines(no_markers, gamevar_mean, type = "l", col = "red")
  arrows(no_markers, PyMSQ_mean - PyMSQ_sd,
         no_markers, PyMSQ_mean + PyMSQ_sd,
         angle = 90, code = 3, col = "blue", length = 0.05)
  arrows(no_markers, gamevar_mean - gamevar_sd,
         no_markers, gamevar_mean + gamevar_sd,
         angle = 90, code = 3, col = "red", length = 0.05)
})
mtext(expression(bold("B")), side = 3, line = 0.5, adj = 0)

# Panel C: Peak Memory Usage vs. Individuals
with(mem_ind_summary, {
  plot(no_individuals, PyMSQ_mean, type = "l", col = "blue",
       ylim = range(c(PyMSQ_mean - PyMSQ_sd, PyMSQ_mean + PyMSQ_sd,
                      gamevar_mean - gamevar_sd, gamevar_mean + gamevar_sd)),
       xlab = "Number of Individuals", ylab = "Peak Memory Usage (GB)",
       axes = FALSE)
  axis(2, col.axis = "black", las = 2)
  box()
  lines(no_individuals, gamevar_mean, type = "l", col = "red")
  arrows(no_individuals, PyMSQ_mean - PyMSQ_sd,
         no_individuals, PyMSQ_mean + PyMSQ_sd,
         angle = 90, code = 3, col = "blue", length = 0.05)
  arrows(no_individuals, gamevar_mean - gamevar_sd,
         no_individuals, gamevar_mean + gamevar_sd,
         angle = 90, code = 3, col = "red", length = 0.05)
  axlab = seq(0, max(no_individuals), length.out=11)
  Axis(side = 1, at = axlab, labels = format(axlab, scientific = FALSE), las = 2)
})
mtext(expression(bold("C")), side = 3, line = 0.5, adj = 0)

# Panel D: Peak Memory Usage vs. Markers
with(mem_mark_summary, {
  plot(no_markers, PyMSQ_mean, type = "l", col = "blue",
       ylim = range(c(PyMSQ_mean - PyMSQ_sd, PyMSQ_mean + PyMSQ_sd,
                      gamevar_mean - gamevar_sd, gamevar_mean + gamevar_sd)),
       xlab = "Number of Markers", ylab = "", axes = FALSE)
  axlab = seq(0, max(no_markers), length.out=11)
  Axis(side = 1, at = axlab, labels = axlab, las = 2)
  axis(4, col.axis = "black", las = 2)
  box()
  lines(no_markers, gamevar_mean, type = "l", col = "red")
  arrows(no_markers, PyMSQ_mean - PyMSQ_sd,
         no_markers, PyMSQ_mean + PyMSQ_sd,
         angle = 90, code = 3, col = "blue", length = 0.05)
  arrows(no_markers, gamevar_mean - gamevar_sd,
         no_markers, gamevar_mean + gamevar_sd,
         angle = 90, code = 3, col = "red", length = 0.05)
})
mtext(expression(bold("D")), side = 3, line = 0.5, adj = 0)

# Add global x and y labels
mtext("Time (min)", at = .75, side = 2, outer = TRUE, cex = 1.2, las = 0, line = 1.8)
```

```r
mtext("Peak memory usage (GB)", at = .25, side = 2, outer = TRUE, cex = 1.2, las = 0, line = 1.8)
mtext("Time (min)", at = .75, side = 4, outer = TRUE, cex = 1.2, las = 0, line = 3.5)
mtext("Peak memory usage (GB)", at = .25, side = 4, outer = TRUE, cex = 1.2, las = 0, line = 3.5)
mtext("Number of individuals", at = .25, side = 1, outer = TRUE, cex = 1.2, las = 0, line = 2)
mtext("Number of markers", at = .75, side = 1, outer = TRUE, cex = 1.2, las = 0, line = 2)

# Common legend
par(fig = c(0, 1, 0, 1), oma = c(0, 0, 0, 0), mar = c(0, 0, 0, 0), new = TRUE)
plot(0, 0, type = 'l', bty = 'n', xaxt = 'n', yaxt = 'n')
legend("bottom", legend = c("gamevar", "PyMSQ"), col = c("red", "blue"),
       lty = c(1, 1), lwd = 1.5, xpd = TRUE, horiz = TRUE, cex = 1.5, seg.len = 1,
       bty = 'n', ncol = 1, y)

dev.off()
```

Now we display Figure 3:

### 4. Performance Comparison Tables

We compare PyMSQ and gamevar using two benchmark datasets:

- Individuals Dataset: Number of markers is fixed at 1000, while the number of individuals varies from 5000 to 100000.
- Markers Dataset: Number of individuals is fixed at 500, while the number of markers varies from 2500 to 50000. Both datasets involve 10 traits and 1 chromosome.

```r
# For the individuals dataset:
time_ind_summary <- perf_ind %>%
  group_by(no_individuals) %>%
  summarise(
    PyMSQ_time = mean(time_PyMSQ, na.rm = TRUE) / 60,
    Gamevar_time = mean(time_gamevar, na.rm = TRUE) / 60
  ) %>%
  mutate(Fold_Faster = Gamevar_time / PyMSQ_time)

mem_ind_summary <- perf_ind %>%
  group_by(no_individuals) %>%
  summarise(
    PyMSQ_memory = mean(peak_memory_usage_PyMSQ, na.rm = TRUE),
    Gamevar_memory = mean(peak_memory_usage_gamevar, na.rm = TRUE)
  ) %>%
  mutate(Memory_Ratio = PyMSQ_memory / Gamevar_memory)

# Merge the summaries for individuals:
individuals_summary <- left_join(time_ind_summary, mem_ind_summary, by = "no_individuals") %>%
  rename(
    `Number of Individuals` = no_individuals,
    `PyMSQ Time (min)` = PyMSQ_time,
    `Gamevar Time (min)` = Gamevar_time,
    `Time Ratio (Gamevar/PyMSQ)` = Fold_Faster,
    `PyMSQ Memory (GB)` = PyMSQ_memory,
    `Gamevar Memory (GB)` = Gamevar_memory,
    `Memory Ratio (PyMSQ/Gamevar)` = Memory_Ratio
  )
```

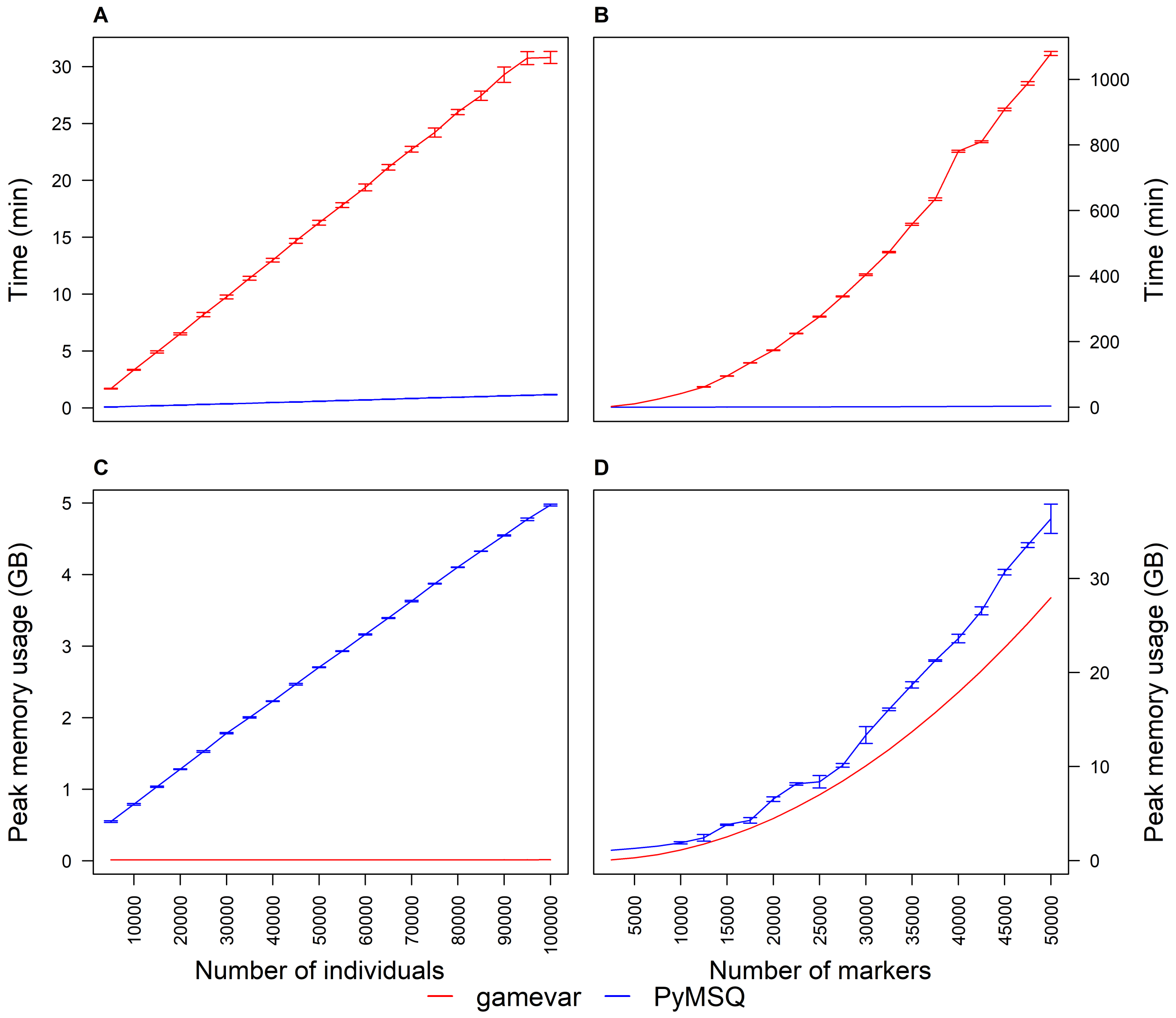


Figure 3: Benchmark plots comparing computation time (Panels A, B) and memory usage (Panels C, D) for PyMSQ and gamevar.

```r
# For the markers dataset:
time_mark_summary <- perf_mark %>%
  group_by(no_markers) %>%
  summarise(
    PyMSQ_time = mean(time_PyMSQ, na.rm = TRUE) / 60,
    Gamevar_time = mean(time_gamevar, na.rm = TRUE) / 60
  ) %>%
  mutate(Fold_Faster = Gamevar_time / PyMSQ_time)

mem_mark_summary <- perf_mark %>%
  group_by(no_markers) %>%
  summarise(
    PyMSQ_memory = mean(peak_memory_usage_PyMSQ, na.rm = TRUE),
    Gamevar_memory = mean(peak_memory_usage_gamevar, na.rm = TRUE)
  ) %>%
  mutate(Memory_Ratio = PyMSQ_memory / Gamevar_memory)

# Merge the summaries for markers:
markers_summary <- left_join(time_mark_summary, mem_mark_summary, by = "no_markers") %>%
  rename(
    `Number of Markers` = no_markers,
    `PyMSQ Time (min)` = PyMSQ_time,
    `Gamevar Time (min)` = Gamevar_time,
    `Time Ratio (Gamevar/PyMSQ)` = Fold_Faster,
    `PyMSQ Memory (GB)` = PyMSQ_memory,
    `Gamevar Memory (GB)` = Gamevar_memory,
    `Memory Ratio (PyMSQ/Gamevar)` = Memory_Ratio
  )
```

**Display the table for the individuals dataset:**

```r
kable(individuals_summary,
      caption = "Performance Comparison (Individuals Dataset)", digits = 4)
```

Table 1: Performance Comparison (Individuals Dataset)

| Number of Individuals | PyMSQ Time (min) | Gamevar Time (min) | Time Ratio (Gamevar/PyMSQ) | PyMSQ Memory (GB) | Gamevar Memory (GB) | Memory Ratio (PyMSQ/Gamevar) |
|---:|---:|---:|---:|---:|---:|---:|
| 5000 | 0.0735 | 1.6917 | 23.0262 | 0.5492 | 0.0139 | 39.5290 |
| 10000 | 0.1340 | 3.3415 | 24.9369 | 0.7896 | 0.0140 | 56.2000 |
| 15000 | 0.1945 | 4.9152 | 25.2685 | 1.0361 | 0.0141 | 73.5025 |
| 20000 | 0.2380 | 6.5075 | 27.3443 | 1.2809 | 0.0141 | 90.6188 |
| 25000 | 0.3052 | 8.1996 | 26.8676 | 1.5272 | 0.0142 | 107.2078 |
| 30000 | 0.3567 | 9.7491 | 27.3340 | 1.7837 | 0.0143 | 125.1062 |
| 35000 | 0.4025 | 11.3988 | 28.3166 | 2.0036 | 0.0143 | 139.8827 |
| 40000 | 0.4731 | 12.9819 | 27.4391 | 2.2311 | 0.0143 | 155.8354 |
| 45000 | 0.5218 | 14.6783 | 28.1301 | 2.4683 | 0.0144 | 171.4445 |
| 50000 | 0.5775 | 16.2707 | 28.1760 | 2.7052 | 0.0144 | 187.2737 |
| 55000 | 0.6475 | 17.8198 | 27.5195 | 2.9305 | 0.0145 | 202.6832 |
| 60000 | 0.6973 | 19.3777 | 27.7896 | 3.1631 | 0.0145 | 217.8085 |
| 65000 | 0.7651 | 21.1444 | 27.6373 | 3.3932 | 0.0145 | 234.1999 |

| Number of Individuals | PyMSQ Time (min) | Gamevar Time (min) | Time Ratio (Gamevar/PyMSQ) | PyMSQ Memory (GB) | Gamevar Memory (GB) | Memory Ratio (PyMSQ/Gamevar) |
|---:|---:|---:|---:|---:|---:|---:|
| 70000 | 0.8220 | 22.7347 | 27.6589 | 3.6307 | 0.0146 | 249.1597 |
| 75000 | 0.8868 | 24.2053 | 27.2962 | 3.8740 | 0.0145 | 266.3079 |
| 80000 | 0.9366 | 26.0018 | 27.7629 | 4.1030 | 0.0146 | 280.3230 |
| 85000 | 0.9942 | 27.4408 | 27.6023 | 4.3259 | 0.0147 | 294.8262 |
| 90000 | 1.0470 | 29.2895 | 27.9747 | 4.5471 | 0.0147 | 309.1979 |
| 95000 | 1.1101 | 30.7444 | 27.6947 | 4.7721 | 0.0147 | 323.7633 |
| 100000 | 1.1609 | 30.7995 | 26.5314 | 4.9705 | 0.0148 | 334.9568 |

**Display the table for the markers dataset:**

```r
kable(markers_summary, caption = "Performance Comparison (Markers Dataset)",
      digits = 4)
```

Table 2: Performance Comparison (Markers Dataset)

| Number of Markers | PyMSQ Time (min) | Gamevar Time (min) | Time Ratio (Gamevar/PyMSQ) | PyMSQ Memory (GB) | Gamevar Memory (GB) | Memory Ratio (PyMSQ/Gamevar) |
|---:|---:|---:|---:|---:|---:|---:|
| 2500 | 0.0333 | 2.4414 | 73.2419 | 1.1052 | 0.0729 | 15.1675 |
| 5000 | 0.0668 | 10.4440 | 156.3479 | 1.2909 | 0.2827 | 4.5664 |
| 7500 | 0.1107 | 24.6128 | 222.4045 | 1.5237 | 0.6321 | 2.4105 |
| 10000 | 0.1708 | 41.5811 | 243.4966 | 1.8872 | 1.1213 | 1.6831 |
| 12500 | 0.2675 | 61.8284 | 231.1198 | 2.4206 | 1.7502 | 1.3830 |
| 15000 | 0.3605 | 95.3007 | 264.3447 | 3.8162 | 2.5188 | 1.5151 |
| 17500 | 0.4679 | 135.2333 | 289.0526 | 4.2710 | 3.4270 | 1.2463 |
| 20000 | 0.5714 | 173.6894 | 303.9894 | 6.5295 | 4.4750 | 1.4591 |
| 22500 | 0.7248 | 224.9720 | 310.3847 | 8.1461 | 5.6627 | 1.4386 |
| 25000 | 0.9020 | 276.2836 | 306.3011 | 8.3819 | 6.9901 | 1.1991 |
| 27500 | 1.0024 | 338.1668 | 337.3515 | 10.1369 | 8.4571 | 1.1986 |
| 30000 | 1.2325 | 404.2205 | 327.9635 | 13.3481 | 10.0639 | 1.3263 |
| 32500 | 1.3395 | 472.9189 | 353.0519 | 16.0859 | 11.8104 | 1.3620 |
| 35000 | 1.6101 | 558.0030 | 346.5678 | 18.6937 | 13.6965 | 1.3648 |
| 37500 | 1.8505 | 634.4724 | 342.8685 | 21.2675 | 15.7224 | 1.3527 |
| 40000 | 2.1511 | 781.0272 | 363.0799 | 23.6134 | 17.8880 | 1.3201 |
| 42500 | 2.3476 | 809.9011 | 344.9935 | 26.5606 | 20.1932 | 1.3153 |
| 45000 | 2.6547 | 908.1433 | 342.0889 | 30.6722 | 22.6381 | 1.3549 |
| 47500 | 3.0119 | 987.5281 | 327.8773 | 33.5492 | 25.2228 | 1.3301 |
| 50000 | 3.2457 | 1078.8873 | 332.4034 | 36.3486 | 27.9472 | 1.3006 |

### 5. Performance Table 3: Benchmark of similarity matrices

Using the same data set for benchmarking Mendelian (co-variability), we benchmarked the time and peak memory usage:

```r
library(dplyr)
library(knitr)


# Read csv files
```

```r
perf_ind <- read.csv("similarity_unsaved.csv")
perf_mark <- read.csv("similarity_mark_unsaved.csv")

# Compute computation time (in minutes) + SD for individuals
time_ind_summary <- perf_ind %>%
  group_by(no_individuals) %>%
  summarise(
    times = mean(time, na.rm = TRUE) / 60,
    time_sd = sd(time, na.rm = TRUE) / 60
  )

mem_ind_summary <- perf_ind %>%
  group_by(no_individuals) %>%
  summarise(
    memory = mean(peak_memory_usage, na.rm = TRUE),
    memory_sd = sd(peak_memory_usage, na.rm = TRUE)
  )

# Markers scenario
time_mark_summary <- perf_mark %>%
  group_by(no_markers) %>%
  summarise(
    times = mean(time, na.rm = TRUE) / 60,
    time_sd = sd(time, na.rm = TRUE) / 60
  )

mem_mark_summary <- perf_mark %>%
  group_by(no_markers) %>%
  summarise(
    memory = mean(peak_memory_usage, na.rm = TRUE),
    memory_sd = sd(peak_memory_usage, na.rm = TRUE)
  )

# Merge time & memory for Individuals
ind_summary <- left_join(time_ind_summary, mem_ind_summary, by = "no_individuals") %>%
  mutate(
    `Time (min)` = paste0(round(times, 2), " ± ", round(time_sd, 2)),
    `Memory (GB)` = paste0(round(memory, 2), " ± ", round(memory_sd, 2))
  ) %>%
  select(no_individuals, `Time (min)`, `Memory (GB)`)

# Merge time & memory for Markers
mark_summary <- left_join(time_mark_summary, mem_mark_summary, by = "no_markers") %>%
  mutate(
    `Time (min)` = paste0(round(times, 2), " ± ", round(time_sd, 2)),
    `Memory (GB)` = paste0(round(memory, 2), " ± ", round(memory_sd, 2))
  ) %>%
  select(no_markers, `Time (min)`, `Memory (GB)`)

# Rename to avoid duplicates
ind_summary_renamed <- ind_summary %>%
  rename(
    "No. Individuals" = no_individuals,
```

```r
    "Time (min) [Ind]" = "Time (min)",
    "Memory (GB) [Ind]" = "Memory (GB)"
  )

mark_summary_renamed <- mark_summary %>%
  rename(
    "No. Markers" = no_markers,
    "Time (min) [Mark]" = "Time (min)",
    "Memory (GB) [Mark]" = "Memory (GB)"
  )

# Combine side by side
results <- cbind(ind_summary_renamed, mark_summary_renamed)

# Print the merged table
kable(results, caption = "Benchmark Results for Similarity Matrix")
```

Table 3: Benchmark Results for Similarity Matrix

| No. Individuals | Time (min) [Ind] | Memory (GB) [Ind] | No. Markers | Time (min) [Mark] | Memory (GB) [Mark] |
|---:|---|---|---:|---|---|
| 5000 | 0.08 ± 0.02 | 0.76 ± 0.05 | 2500 | 0.04 ± 0.01 | 0.37 ± 0.03 |
| 10000 | 0.17 ± 0.01 | 1.36 ± 0.04 | 5000 | 0.07 ± 0 | 0.63 ± 0.03 |
| 15000 | 0.3 ± 0 | 2.33 ± 0.16 | 7500 | 0.1 ± 0 | 1.04 ± 0.06 |
| 20000 | 0.44 ± 0 | 3.33 ± 0.17 | 10000 | 0.17 ± 0 | 1.57 ± 0.19 |
| 25000 | 0.66 ± 0.02 | 4.93 ± 0.32 | 12500 | 0.23 ± 0 | 2.33 ± 0.06 |
| 30000 | 0.89 ± 0.02 | 6.78 ± 0.32 | 15000 | 0.34 ± 0 | 3.15 ± 0.05 |
| 35000 | 1.16 ± 0.02 | 9.12 ± 0.56 | 17500 | 0.4 ± 0.01 | 4.16 ± 0.04 |
| 40000 | 1.45 ± 0.01 | 11.71 ± 0.77 | 20000 | 0.53 ± 0 | 5.93 ± 0.22 |
| 45000 | 1.82 ± 0.02 | 14.32 ± 0.62 | 22500 | 0.64 ± 0 | 7.66 ± 0.12 |
| 50000 | 2.2 ± 0 | 16.76 ± 0.29 | 25000 | 0.79 ± 0.02 | 7.92 ± 0.03 |
| 55000 | 2.62 ± 0 | 21.18 ± 0.58 | 27500 | 0.94 ± 0 | 9.45 ± 0.04 |
| 60000 | 3.06 ± 0.02 | 25.8 ± 0.82 | 30000 | 1.07 ± 0 | 12.24 ± 0.41 |
| 65000 | 3.56 ± 0 | 29.27 ± 0.9 | 32500 | 1.24 ± 0.01 | 15.05 ± 0.36 |
| 70000 | 4.07 ± 0.02 | 33.67 ± 0.43 | 35000 | 1.44 ± 0 | 17.57 ± 0.28 |
| 75000 | 4.6 ± 0.03 | 37.55 ± 0.64 | 37500 | 1.67 ± 0.1 | 20.09 ± 0.19 |
| 80000 | 5.26 ± 0.03 | 44.73 ± 1.62 | 40000 | 1.89 ± 0.08 | 22.5 ± 0.38 |
| 85000 | 5.86 ± 0.05 | 50.46 ± 0.62 | 42500 | 2.07 ± 0.06 | 24.78 ± 0.68 |
| 90000 | 6.48 ± 0.05 | 55.42 ± 0.52 | 45000 | 2.32 ± 0.07 | 30.03 ± 0.43 |
| 95000 | 7.15 ± 0.07 | 60.44 ± 0.67 | 47500 | 2.54 ± 0 | 32.82 ± 0.35 |
| 100000 | 7.93 ± 0.09 | 69.58 ± 1.87 | 50000 | 2.8 ± 0.02 | 34.28 ± 0.22 |
